# Perceiving and remembering speech depend on multifractal nonlinearity in movements producing and exploring speech

**DOI:** 10.1101/2021.03.28.437328

**Authors:** Lauren Bloomfield, Elizabeth Lane, Madhur Mangalam, Damian G. Kelty-Stephen

**Author notes:** Author for correspondence: Damian G. Kelty-Stephen. **Ethics statement.** All participants provided informed consent approved by the Institutional Review Board (IRB) at Grinnell College, Grinnell, IA. **Data accessibility.** All data analyzed in the present study are available as supplementary material. **Author contributions.** L.B., E.L., and D.G.K-S. conceived and designed research; L.B., E.L., and D.G.K-S. performed experiments; L.B., E.L., M.M., and D.G.K-S. curated and analyzed data; L.B., E.L., M.M., and D.G.K-S. interpreted results of experiments; M.M. and D.G.K-S. prepared figures; L.B., E.L., M.M., and D.G.K-S. drafted manuscript; L.B., E.L., M.M., and D.G.K-S. edited and revised manuscript; L.B., E.L., M.M., and D.G.K-S. approved final version of manuscript. **Competing interests.** The authors have no competing interests to declare.

## Abstract

Speech perception and memory for speech require active engagement. Gestural theories have emphasized mainly the effect of speaker’s movements on speech perception. They fail to address the effects of listener movement, focusing on communication as a boundary condition constraining movement among interlocutors. The present work attempts to break new ground by using multifractal geometry of physical movement as a common currency for supporting both sides of the speaker-listener dyads. Participants self-paced their listening to a narrative, after which they completed a test of memory querying their narrative comprehension and their ability to recognize words from the story. The multifractal evidence of nonlinear interactions across timescales predicted the fluency of speech perception. Self-pacing movements that enabled listeners to control the presentation of speech sounds constituted a rich exploratory process. The multifractal nonlinearity of this exploration supported several aspects of memory for the perceived spoken language. These findings extend the role of multifractal geometry in the speaker’s movements to the narrative case of speech perception. In addition to posing novel basic research questions, these findings make a compelling case for calibrating multifractal structure in text-to-speech synthesizers for better perception and memory of speech.

## 1. Introduction

Listening to speech entails active engagement. Even when listening is most passive (e.g., when listening to a podcast on headphones), beyond vibrations from the eardrums, the listener consults the lexicon, a trove of possible words and their corresponding meaning [1]. The listener must explore the characteristics of the sounds themselves. Listeners can potentially face an uncertain terrain when exploring the speech around them. The present study investigates how we may use movement to explore speech that supports perception of and cognition about spoken words.

Speech perception depends on the articulatory motor processes shaping speech’s acoustic features [2–4]. Motor processes reshape phonemes in context- and sequence-dependent ways (e.g., in coarticulation) that text-to-speech synthesis has struggled to emulate [5]. Speech sounds reflect movements beyond articulators to distal parts of the body [6]—potentially explaining their utility for diagnosing Parkinson’s disease [7]. However, movement is now being recognized as an aspect of speech that otherwise passive listeners can use [8]. Gestural theories have been at most ambivalent about the role of listener’s actions in perception and cognition of spoken words (e.g., compare [9] with [10,11]), most recently focusing on communication as a boundary condition constraining movement among interlocutors [12]. However, perceiving speech becomes more difficult when we listen to movement systems that act less like we do ourselves, as when we listen to a speaker from another linguistic group [13].

Here, we test an experimental contrast of a human speaker with the text-to-speech synthesis in a between-groups design. Text-to-speech synthesizers may lack many context- and sequence-dependencies in human speech [14], and they dramatically reshape the movement-structured terrain for the listener. For present purposes, we use the speech waveform as an operationalization of speech movements. Although it is not an anatomically specific measurement of individual speech articulators, the speech waveform carries the imprint of fine kinetic and kinematic markers of speech articulators [15], subtle enough to aid in distinguishing phonation type [16], misarticulated stops from cleft lip or palate [17], and dysarthrias [18].

The geometry of listener movements for exploring speech sounds may be no less crucial than speaker movements. Auditory perception has strong roots in a listener’s bodywide capability for physical movement [19–25], and spoken-language perception depends on movement, e.g., with activity of the motor cortex in covert imitation of speech sounds supporting interactive alignment across many scales of dyadic conversation [26]. For instance, even in the case of listening to recorded speech, the spatial organization of speech in the space around the listener’s body facilitates spoken-word perception under difficult listening circumstances, for example, with noise [27] or reverberation [28]. We can elicit overt exploratory behaviors, such as key pressing, during speech perception through the “self-paced listening” paradigm requiring participants to make key presses to signal readiness for consecutive words in a sequence [29]. Self-pacing affords the reader more control in exploring linguistic stimuli, with progress or pause indicating the waxing and waning of focus and processing, and giving readers with attentional lapses or sensory deficits a greater opportunity to allocate attention and memory for speech stimuli [30,31].

We align articulatory movement for speech production reflected by the speech waveform with exploratory self-pacing movements for speech perception within a single geometric formalism: multifractal geometry. Multifractal geometry may reveal a causal relationship between (1) motor processes underlying speech stimuli and (2) listener movements for exploring those stimuli. The multifractal structure might predict how well individual listeners process the spoken-word stimulus (figure 1*a*, top right) and later exhibit their memory for the spoken words (figure 1*a*, bottom left). In Part-1, participants listened to a story, word-by-word, articulated by either human speech or text-to-speech synthesizer. This manipulation allowed variations in speech-stream geometry across different words and movement systems (i.e., natural human speech vs. computer-simulated analog). In Part-2, participants completed a memory test querying their comprehension of and memory for words from the story.

**Figure 1.**
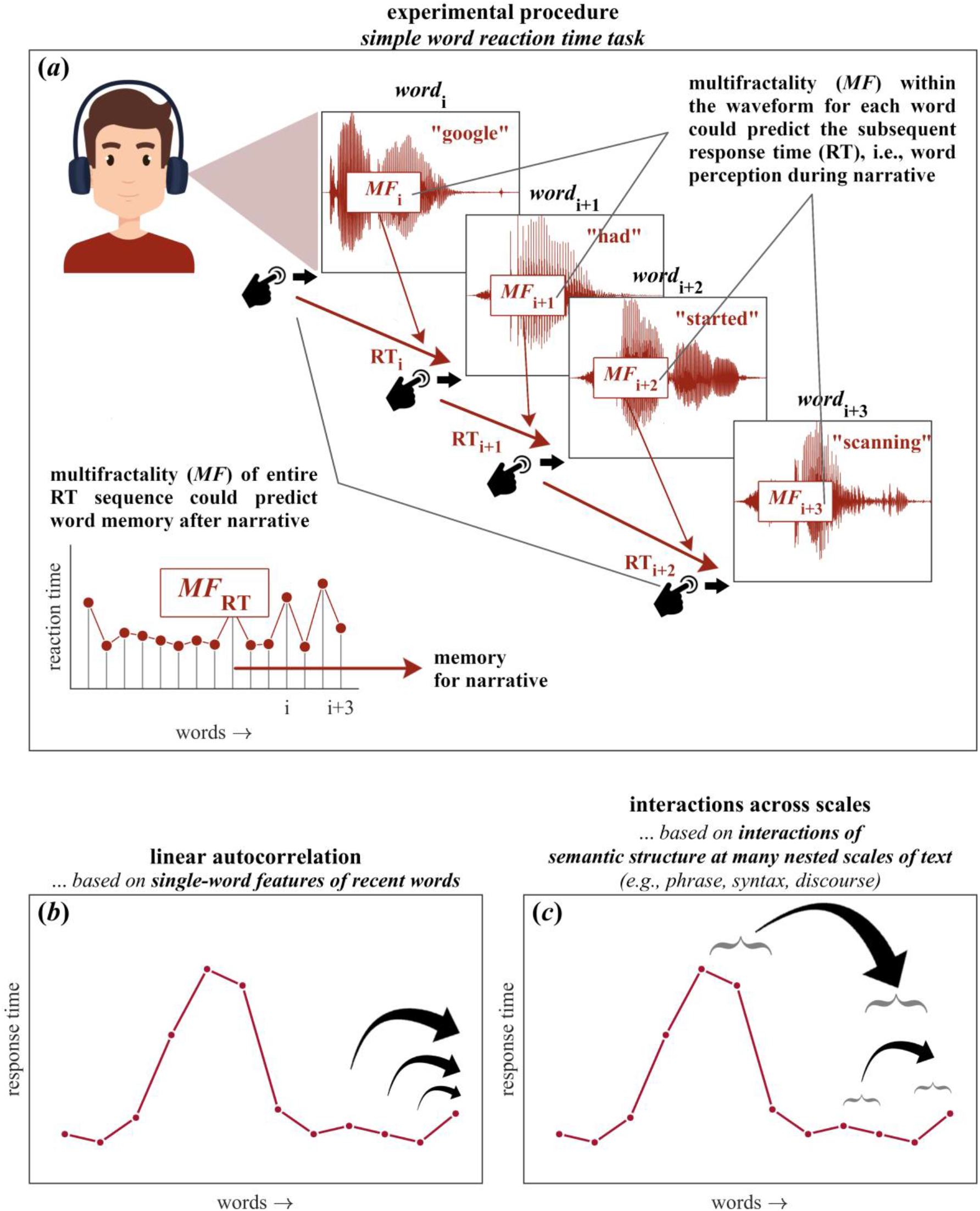
Experimental procedure and two perspectives response times (RT). (*a*) Each word presented to a participant through headphones elicited a button press when the participant was ready for the next word. The multifractal and psycholinguistic properties of each word were used to predict response time to test Hypothesis-1. Multifractal attributes (and standard descriptive statistics) for each Hypothesis-2: whether multifractal attributes of the RT series predicted memory and comprehension of the story (see Section 2.2 for details). (*b*) The autoregressive perspective that each word’s RT entails the cumulative summing of subsequent and independent processes regarding lexical aspects of preceding words, with longer arrows indicating effects of previous behavior at independent, non-overlapping time lags. (*c*) The multifractal perspective that each word’s RT entails the interactions unfolding among processes across many nested time scales of text (e.g., word, phrase, and discourse), with arrows indicating the effects of previous behavior at different time lags. Horizontal braces schematize how effects of previous behavior at longer, coarser scales (top of panel c) may influence shorter, finer scales.

### 1.1. Multifractal geometry helps to formalize the movement-based support for conveying information

Multifractal geometry is a generalization of fractal geometry that encompasses ‘multiple’ degrees of fractal structure [32–34]. Rare in psycholinguistic research (e.g., [35–37]), variation in fractal structure has begun to demonstrate usefulness in modeling gestural supports for perceiving speech [3,4] as well as modeling how readers explore language in a narrative [38].

### 1.2. Multifractality offers statistical estimation of nonlinear interactions across scales, indicating how systems blend current, fleeting behaviors into longer-term behaviors

Multifractal analysis examines how variation depends on time in a measured behavior and how uniformly or unevenly variation unfolds from a brief glimpse to a longer span. Multifractal geometry evaluates variability through a variety of scales within a single measurement. For instance, the analysis segments a time-series measurement, such as audio waveform, into sequential but nonoverlapping bins of the same small size. Each bin represents a short subset of the whole movement at one short time scale. The analysis then estimates variability across those different bins of the same size. Then, multifractal geometry iterates this process for progressively longer timescales.

The average amount of variability in measurement grows as a power law of timescale. Only one power law relates average proportion to timescale for homogeneous time series in which brief and long-term events do not interact. However, for heterogeneous measurements of systems with interactions across scales, a single measurement can contain multiple proportion-to-time power laws. The word ‘multifractal’ reflects that heterogeneous systems can exhibit multiple power-law relationships each with different fractional exponents. This variety of fractional power-law exponents constitutes a ‘multifractal spectrum.’

Two multifractal statistics indicate the strength of relationships propagating brief events across longer timescales. The first metric, the multifractal spectrum width *Δα*, is conceptually analogous to standard deviation but reflects a mixture of linear variability and nonlinear variability. The linear variability concerns histogram width and average drift over time. But the nonlinear variability crucially concerns interactions across scales. Another metric ‘multifractal nonlinearity’ *t*_MF_ is a *t*-statistic indicating difference of *Δα* from a sample of expected *Δα* values from strictly linear models of the measurement [39]. Bottom panels of figures 2*b* and 2*c* schematize this difference between linear structure and nonlinear interactions across timescales.

**Figure 2.**
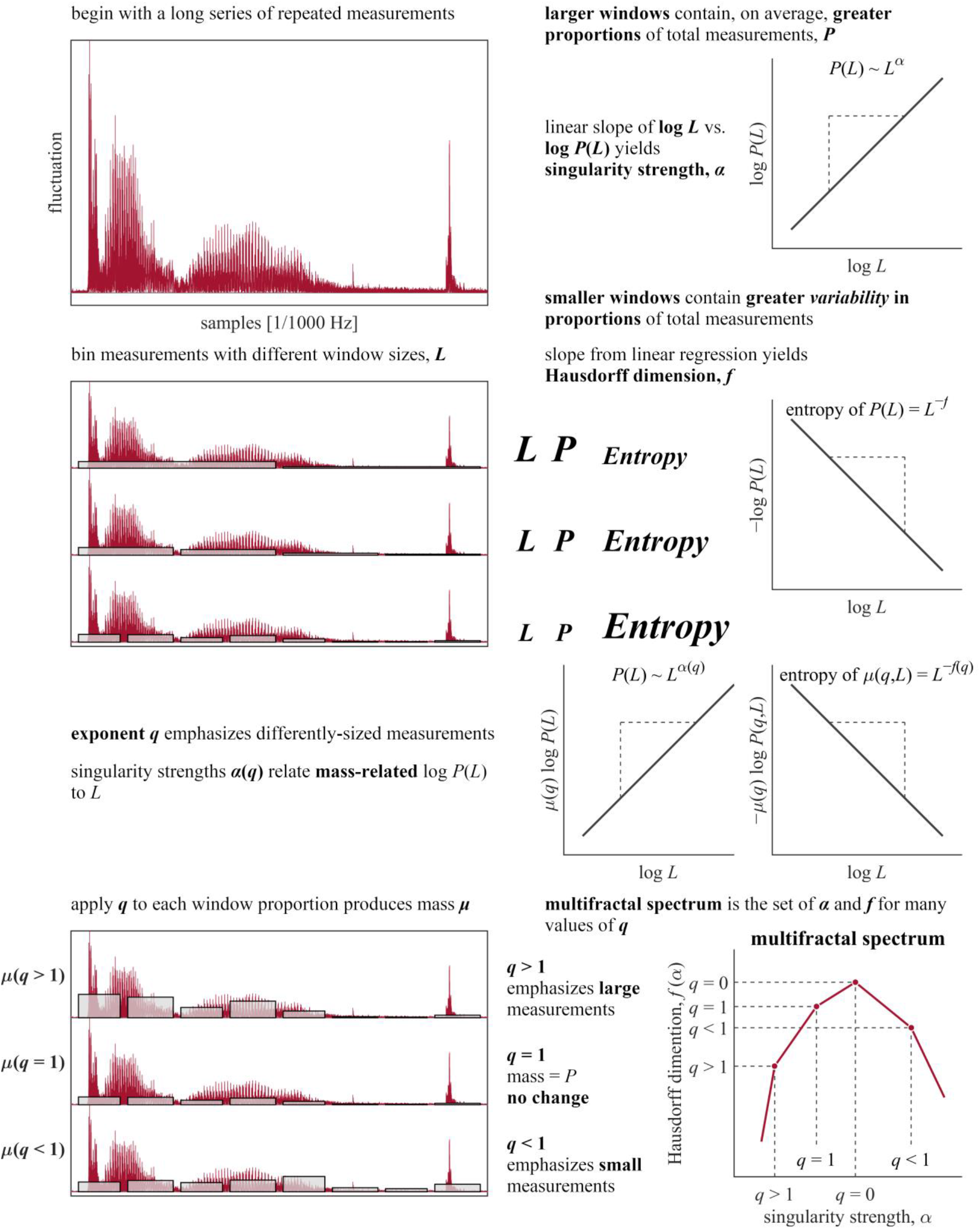
Chhabra and Jensen’s [66] method used to estimate multifractal spectrum *f*(*α*). The method evaluates how well a measured series (top left) fits a general expectation about relationships between proportion and the length of observation. This procedure tiles the series with nonoverlapping windows of progressively greater length (2^nd^ row left). Windowed subsets will cover greater proportion of total series as length increases (top right), with lesser Shannon entropy (2^nd^ row right). Smaller and larger values of a continuous q exponent emphasize sparser and denser portions of the series, respectively (3^rd^ row left). Applying q to window proportions produces a “mass” quantity (bottom left). “Mass”-weighted proportion and the Shannon entropy of “mass” both exhibit scaling relationships with window length (3^rd^ row right). Scaling exponents for these relationships at each q value specify an asymmetric inverted-U shaped curve known as the multifractal (or singularity) spectrum. See Section 2.4.1 for further details.

Multifractality is the natural, logical outcome when different timescales interact. Models of speech production/perception include hierarchical models that nest fine, relatively “analog” details of movement and stimulation into progressively broader and more abstract neuroscientific and linguistic categories, structures, or expectations. These models often manifest in Bayesian linear models [40] or convolutional neural networks [41]. These elegant models fall short of explaining the speech-production/perception outcomes in evidence [42]. Their hierarchical structure is so static that as to force symmetrical repetition and mimicry sooner than generating dynamic and time-evolving synergistic structures [43]. The use of recurrence quantification has been a significant step towards examining language use across multiple scales without strictly linear assumptions (e.g., [43]). However, multifractal modeling can address more specifically the role of nonlinearity in cross-scale interactions in producing the asymmetric waxing or waning of specific synergistic coordinations. So, multifractality may answer current calls for formalisms capable of developing falsifiable hypotheses about synergies in language use [13].

Both speech production and language perception exemplify just such nonlinear interactions across timescales. Longer-term speech sequences reshape the brief articulation of a single syllable [5]. Communicative sounds exhibit a range of fractional power-law exponents [44]. Multifractality nonlinearity *t*_MF_ may predict expressiveness [45]. Speech perception adjusts to compensate for coarticulatory interaction across time [2] and tailors these compensations over longer timescales [46]. Indeed, research has already modeled the effects of the multifractal structure of speech waveforms in supporting speech perception [3,4]. We choose to use the waveform rather than an auditory envelope. Envelopes involve smoothing procedures that may serve multifractal analysis for very long series (e.g., whole bird song phrases [45]). We chose not to use an envelope of any sort because we risked removing finer vibrations in these briefer single-word recordings that would support multifractal modeling over a wider range of scales.

### 1.3. Multifractal fluctuations in measurements of movement predict how movements support the pick-up, coordination, and later use of information for perceptual judgments

Multifractal geometry has repeatedly captured predictive information about how movement systems loosen their constraints to absorb and then propagate perceptual information. For example, this predictive capacity can be seen in haptic [47,48] and visual perception [49,50], as well as how movement systems integrate information across the body [51,52] and across different modalities [53]. In psycholinguistic domains, greater multifractal nonlinearity in self-paced reading may keep readers’ expectations loose and open-minded enough to navigate unexpected plot twists more fluently, (i.e., with a lower increase in subsequent reading-time), indicating a lighter processing load [38].

Multifractal fluctuations support the sharing of information necessary for dexterous behavior across organisms [54,55], just like they do within organisms. Indeed, attention to a speaker engenders a fractal-like ‘complexity matching’ supporting the listener’s comprehension of linguistic content in the signal [56–58]. There has been growing curiosity in how these findings relate to insights from the multifractal formalism [58,59]. We now test whether such multifractal geometry supports this social-information exchange in speech production and perception.

### 1.4. Two hypotheses: Multifractality nonlinearity supports the uptake and memory for spoken-language

#### 1.4.1. Hypothesis-1: Greater multifractal nonlinearity of speech sounds predicts more fluent response to individual words, prolonging fluency of perceiving text-to-speech synthesized words comparable to human-voice words

We used a self-paced listening task to experimentally test the effect of variations in multifractality—both from word to word within the same speaker and between two speakers with different styles of producing speech. We enlisted a human speaker and a text-to-speech synthesizer to pronounce each word of a narrative separately but in sequence. Despite supporting students with difficulty reading by sight [60], the speech-production in text-to-speech is unfamiliar and perceptually challenging [61]. Because of the potential role of multifractal nonlinearity in supporting expressivity, we predicted that multifractal nonlinearity in speech would speed the word-by-word response in self-paced listening (Hypothesis-1a). Likewise, we predicted that multifractal nonlinearity would support the fluency of text-to-speech synthesis perception and defer any gradual slowing of the self-pacing in response to text-to-speech speech relative to human speech (Hypothesis-1b). We explored effects of raw multifractal spectrum width without anticipating any single direction.

#### 1.4.2. Hypothesis-2: Greater multifractal spectrum width and greater nonlinearity of the self-paced key press interval time series would predict stronger performance on post-narrative test of comprehension and recognition memory

We predicted that greater multifractal nonlinearity across key-press sequences would predict better performance on a post-narrative memory test. We explored effects of raw multifractal spectrum width without anticipating any single direction.

## 2. Materials and methods

### 2.1. Participants

Nine healthy men and eleven healthy women (*mean*±*s.d.* age, 20.10±1.29 years) participated after providing informed consent. Three participants reported having a learning disability, two with attention deficit hyperactivity disorder (ADHD) and 1 with dyslexia, amounting to the same proportion of people with the same diagnoses in the United States population [62,63].

### 2.2. Experimental task and procedure

Participants were randomly assigned to hear the voice of either an adult woman or Acapela’s U.S. English text-to-speech female voice ‘Sharon’ (Acapela Inc., Mons, Belgium) using iPad app ‘Voice Dream.’ Both produced speech recordings of 2,027 words in sequence from *The Atlantic* article “Torching the Modern-Day Library of Alexandria.” The text-to-speech recording had two fewer words because of a text-parsing error (e.g., omitting to convert “$” symbols to the word “dollars” after the amount). Human speech and text-to-speech both produced words interspersed with pauses to allow parsing. Words were not controlled to have the same pitch and duration between speakers, as the primary feature of interest was the multifractal nonlinearity of the speaker’s word production. Hence, the within-speaker assessment of nonlinearity was the higher priority and controlled for average pitch and duration effects through comparison to surrogates built to mimic the amplitude spectrum and duration. Multiple standard psycholinguistic features of these words were assessed: word frequency, ‘logWF,’ in SUBTLEX [64]; and number of orthographic and phonological neighbors—‘OrthN’ and ‘PhonN,’ respectively— in CLEARPOND [65].

E-Prime software (Psychology Software Tools Inc., Pittsburgh, PA) presented individual audio recordings of each word in original sequence through headphones. Participants sat at an E-Prime-ready computer and were instructed: “Listen to the audio stimuli and press the spacebar after you feel as though you have understood the word you just heard. Try to pay attention to the passage because comprehension and word-memory questions will be asked at the end of the experiment. However, if you miss a word, do not worry and continue to move on because you cannot go back.” Word recordings played only once immediately as the spacebar was released. Participants could press the spacebar again, either while or after the recording played, to hear the next word, potentially interrupting the full word before its completion in order to hear the next word. E-Prime recorded the response time (RT) in milliseconds (up to one decimal place) from each word onset to subsequent button press. After the completion of the self-paced listening task, participants completed a 20-item pen-and-paper test (see Reading Comprehension Test in Supplementary Note S1). The first ten items prompted participants to indicate whether each of the ten sentences was ‘true’ or ‘false’ based on their comprehension of the narrative. The later ten items prompted participants to demonstrate word-recognition memory by indicating whether each of 10 words appeared in the text. Only five of the words appeared in the text; we made seven of the sentences true so as not to give participants the chance to guess that both tests should have the same number of incorrect items.

### 2.3. Multifractal metrics

We computed multifractal metrics for the absolute value of each spoken word’s audio waveform (whether from human speech or text-to-speech) and each participant’s RT sequence. The following summary of the metrics is brief and mathematically abbreviated. We invite readers interested in a longer, more descriptive explanation to examine an open-access resource here [66] for both a more conversational introduction and R syntax for estimating these metrics.

#### 2.3.1. Multifractal spectra width

Chhabra and Jensen’s direct method [67] samples non-negative series *u*(*t*) at progressively larger scales such that proportion of signal *P*_*i*_(*L*) falling within the *i*^*th*^ bin of scale *L* is

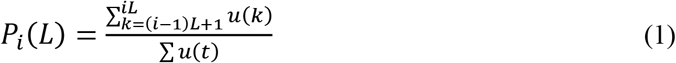

As *L* increases, *P*_*i*_(*L*) represents progressively larger proportion of *u*(*t*),

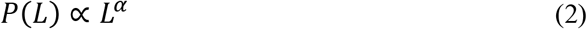

suggesting growth of proportion according to ‘singularity’ strength *α*. *P*(*L*) exhibits multifractal dynamics when it grows heterogeneously across timescales *L* according to multiple potential fractional singularity strengths, such that

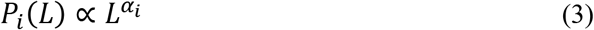

whereby each *i*^*th*^ bin may show a distinct relationship of *P*(*L*) with *L*. The spectrum of singularities is itself the multifractal spectrum, and its width *Δα*(*α*_*max*_ − *α*_*min*_) indicates the heterogeneity of these relationships between proportion and timescale [32,68].

This method estimates *P*(*L*) for *N*_*L*_ nonoverlapping bins of *L*-sizes and accentuates higher or lower *P*(*L*) by using parameter *q* > 1 and *q* < 1, respectively, as follows

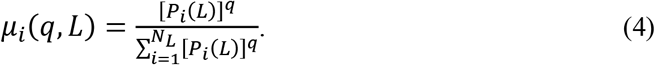

See *figure* 2.

*α*(*q*) is the singularity for *μ*(*q*)-weighted *P*(*L*) estimated by

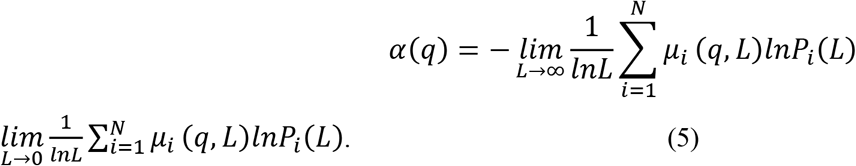

Estimates *α*(*q*) belong to the multifractal spectrum if Shannon entropy of *μ*(*q*, *l*) scales with *L* according to a dimension *f*(*q*), where

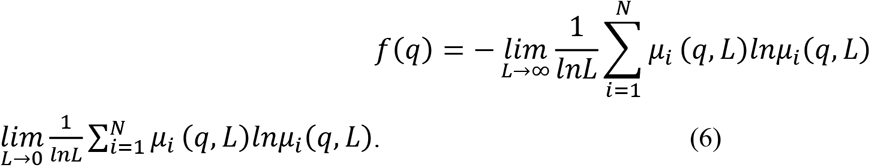

For *q* generating scaling relationships (Eqs. 5 and 6) with correlation coefficient, *r* > 0.95, the parametric curve (*α*(*q*), *f*(*q*)) or (*α*, *f*(*α*)) constitutes the multifractal spectrum with width *Δα* = *α*_*max*_ − *α*_*min*_ (figure 2, bottom right).

#### 2.3.2. Multifractal nonlinearity

To estimate how much observed multifractality reflects nonlinearity [69], *Δα* of each original series was compared to *Δα* of 32 surrogate series obtained using Iterated Amplitude Adjusted Fourier Transformation (IAAFT) [39]. IAAFT generates surrogates that randomize phase ordering of the series’ spectral amplitudes while preserving only linear temporal correlations. We defined multifractal nonlinearity as the one-sample *t*-statistic (henceforth, *t*_MF_) comparing *Δα* of the original series to that of the surrogates (figure 3).

**Figure 3.**
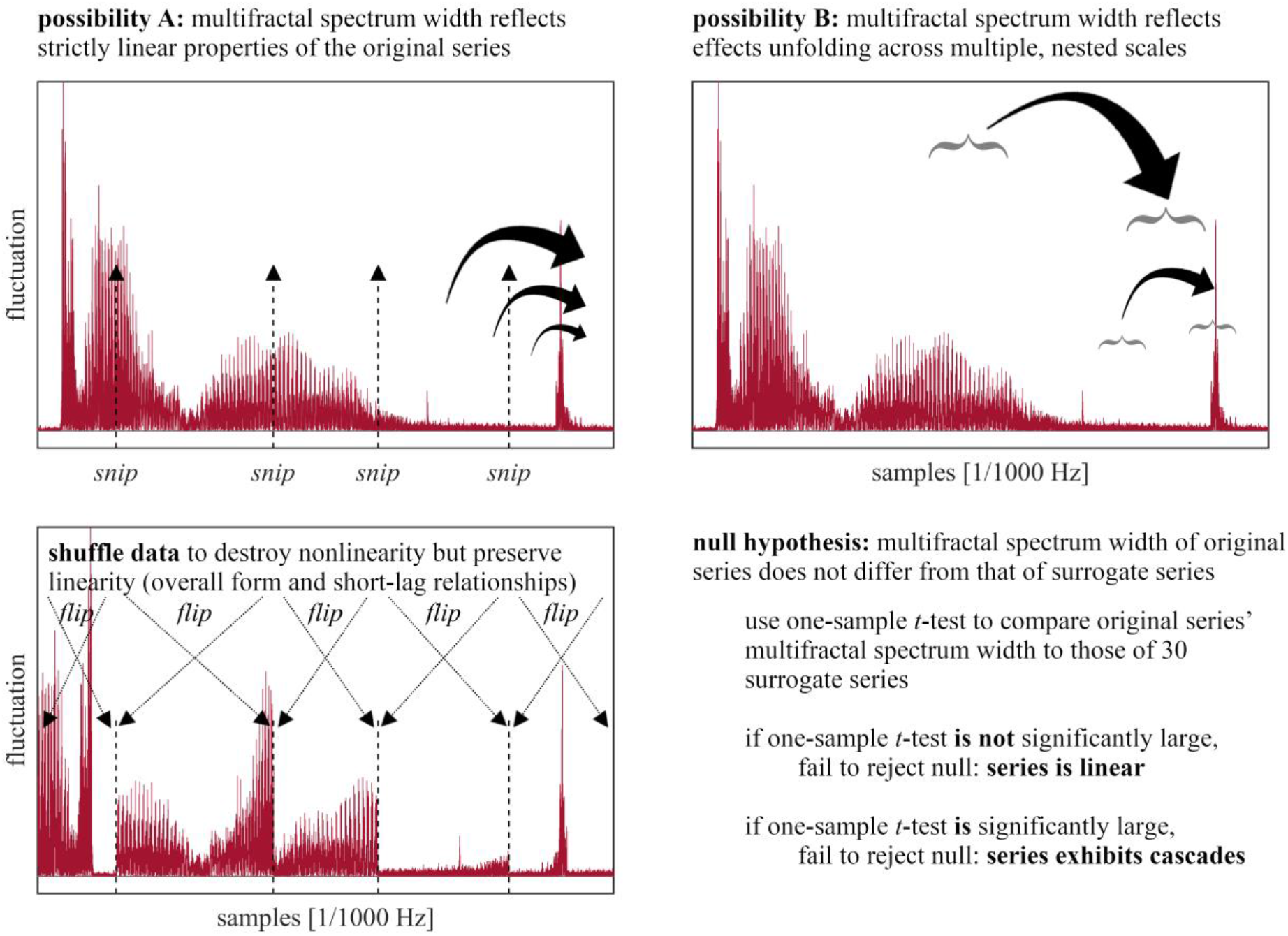
Surrogate analysis used to identify whether a series exhibits multifractality due to nonlinearity (cascades). Speech waveforms may reflect effects unfolding over independent timescales (top left) or interacting timescales (top right). IAAFT procedure preserving linear structure while destroying the original sequence and so any potential interactions across scale. Multifractal-spectrum widths differences between original and surrogate series offers indicates strength of interactions across timescale. See Section 2.4.2 for further details.

### 2.4. Statistical analysis

To test Hypothesis-1, a Poisson regression of self-paced word RT using glmer() in the package *lme4* [70] for *R* [71] included effects of logarithmic sound length (lnSL), word frequency (logWF), number of phonological neighbors (PhonN), number of orthographic neighbors (OrthoN), text-to-speech (TTS = 1 and 0 for synthesizer and human speech, respectively), and *t*_MF_. It also included two time predictors spanning trials across the experiment: word# (word number incrementing by 1 with each word) and, because RT shows a general power-law decrease across progressive words for self-paced processing of narrative text [72], 1/word#. The model included the interaction effects of TTS×word#, as well the interactions 1/word#×TTS×logWF, 1/word#×TTS×PhonN, 1/word#×TTS×OrthN, 1/word#×TTS×*t*_MF_, in addition to all other lower-order interactions. Additional terms or interactions either were nonsignificant or led to poor model convergence (e.g., Δ*α*). We used Poisson regression because E-Prime records RT in discrete units (i.e., tenths of a millisecond). Poisson regression is regularly used to model similarly discrete time-to-events [73–77].

To test Hypothesis-2, a Poisson regression of cumulative correct answers in the post-narrative test included effects of test type (WRM = word recognition memory as opposed to comprehension), condition (TTS = having heard text-to-speech versus human speech), orthogonal cubic polynomials of questions 1 through 10 (calculated using the *R*-function poly() [71]) and various metrics on individual participants’ self-paced RT series including linear features (mean and median) and multifractal metrics (Δ*α* and *t*_MF_). Modeling tested typeWRM×conditionTTS×RT-Δ*α*, typeWRM×conditionTTS×RT-*t*_MF_, conditionTTS×log(RT-mean), conditionTTS×log(RT-median), as well as question×typeWRM and question×Δ*α*, and all lower-order interactions. This model also included a dichotomous predictor ‘LD-status’ (1 or 0 = yes or no) to control any benefits that a learning disability might experience due to the self-paced format [30,31] or their greater fluency with text-to-speech synthesis [78]. Note that LD-status failed to improve the model for Hypothesis 1 and so was omitted from that model. The pattern of significant effects were the same with or without LD-status.

Both models used 1,000 resamplings to generate bootstrapped standard errors, *p*-values, and power estimates. All significant predictors showed at least 80% statistical power (i.e., > 800 resamplings yielded nonzero coefficients in the same direction as the original data) [79].

## 3. Results

Figure 4 shows illustrative output from multifractal analysis of both speech and RT sequences.

**Figure 4.**
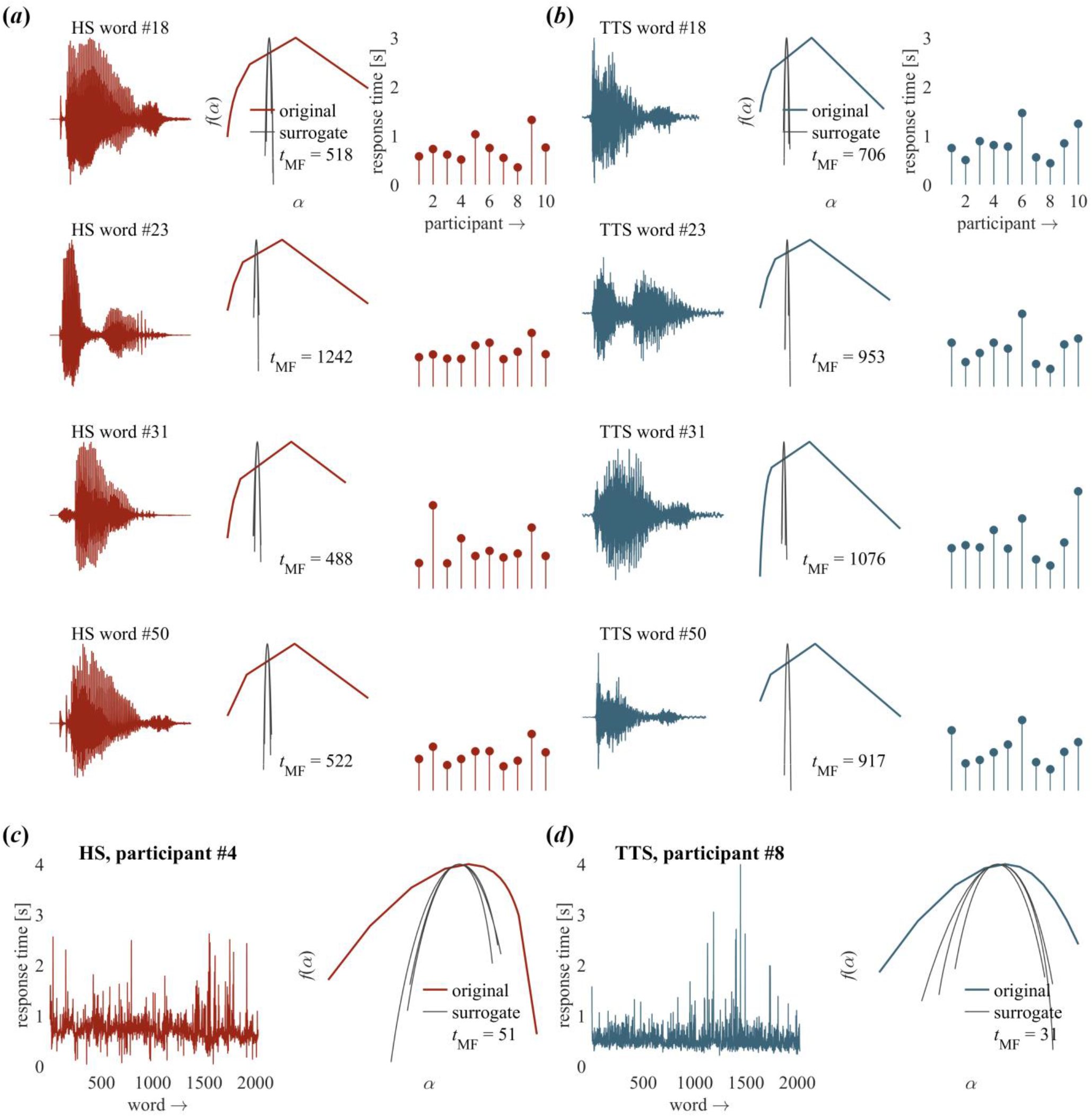
Multifractal spectra of representative stimuli and response times (RTs). Human-speech (a) and text-to-speech (b) words corresponding response times (RTs) for each participant (*n* = 10 in each condition). Multifractal spectrum of RTs for a representative participant hearing human speech (*c*) and for a representative participant hearing text-to-speech condition (*d*).

### 3.1. Model of word-by-word RT

RT for human speech (*mean* = 599.46, *s.e.m.* = 2.17) was lower than for text-to-speech (*mean* = 654.80, *s.e.m.* = 2.44). Table 1 lists this model’s coefficients.

**Table 1.** Coefficients of the Poisson regression of self-paced word RT^a^ (section 3.1).

| effect<sup>b</sup> | *mean* | *s.e.m.*<br>(bootstrapped) | *p*<sup>c</sup><br>(bootstrapped) |
| --- | --- | --- | --- |
| intercept | $5.54 \times 10^0$ | $2.53 \times 10^{-1}$ | **$6.22 \times 10^{-107}$** |
| lnSL | $3.44 \times 10^{-1}$ | $2.42 \times 10^{-2}$ | **$2.83 \times 10^{-46}$** |
| logWF | $-3.22 \times 10^{-2}$ | $2.60 \times 10^{-3}$ | **$2.49 \times 10^{-35}$** |
| PhonN | $-3.97 \times 10^{-4}$ | $2.06 \times 10^{-4}$ | **$2.73 \times 10^{-02}$** |
| OrthN | $-8.26 \times 10^{-5}$ | $4.88 \times 10^{-4}$ | $4.33 \times 10^{-1}$ |
| TTS | $-7.16 \times 10^{-2}$ | $1.75 \times 10^{-2}$ | **$2.19 \times 10^{-5}$** |
| $t_{MF}$ | $-2.77 \times 10^{-5}$ | $1.06 \times 10^{-5}$ | **$4.60 \times 10^{-3}$** |
| TTS×logWF | $1.65 \times 10^{-2}$ | $3.65 \times 10^{-3}$ | **$3.29 \times 10^{-6}$** |
| TTS×PhonN | $5.66 \times 10^{-5}$ | $3.48 \times 10^{-4}$ | $4.35 \times 10^{-1}$ |
| TTS×OrthN | $7.42 \times 10^{-4}$ | $7.53 \times 10^{-4}$ | $1.62 \times 10^{-1}$ |
| TTS$\times t_{MF}$ | $3.46 \times 10^{-5}$ | $1.66 \times 10^{-5}$ | **$1.87 \times 10^{-2}$** |
| word# | $-2.66 \times 10^{-4}$ | $5.44 \times 10^{-6}$ | $0.00 \times 10^0$ |
| TTS×word | $1.42 \times 10^{-4}$ | $8.04 \times 10^{-6}$ | **$2.55 \times 10^{-70}$** |
| 1/word# | $1.02 \times 10^0$ | $3.20 \times 10^{-1}$ | **$7.21 \times 10^{-4}$** |
| 1/word#×logWF | $2.84 \times 10^{-1}$ | $2.60 \times 10^{-3}$ | **$2.49 \times 10^{-35}$** |
| 1/word#×PhonN | $1.73 \times 10^{-2}$ | $1.96 \times 10^{-2}$ | $1.89 \times 10^{-1}$ |
| 1/word#×OrthN | $-8.35 \times 10^{-2}$ | $5.83 \times 10^{-2}$ | $7.61 \times 10^{-2}$ |
| 1/word#×TTS | $-4.06 \times 10^{-1}$ | $5.29 \times 10^{-1}$ | $2.21 \times 10^{-1}$ |
| 1/word#$\times t_{MF}$ | $-1.43 \times 10^{-4}$ | $5.21 \times 10^{-4}$ | $3.92 \times 10^{-1}$ |
| 1/word#×TTS×logWF | $-8.41 \times 10^{-2}$ | $1.97 \times 10^{-1}$ | $3.35 \times 10^{-1}$ |
| 1/word#×TTS×PhonN | $8.21 \times 10^{-2}$ | $2.59 \times 10^{-2}$ | **$7.65 \times 10^{-4}$** |
| 1/word#×TTS×OrthN | $3.03 \times 10^{-2}$ | $8.40 \times 10^{-2}$ | $3.59 \times 10^{-1}$ |
| 1/word#×TTS$\times t_{MF}$ | $1.81 \times 10^{-3}$ | $6.47 \times 10^{-4}$ | **$2.55 \times 10^{-3}$** |

<sup>a</sup>model in R syntax: `glmer(I(10*RT) ~ lnSL + I(1/word#)*logWF*condition + I(1/word#)*PhonN*condition + I(1/word#)*OrthN*condition + I(1/word#)*condition*tMF + word#*condition + (1|participant), data = DATA, family = poisson).`
<sup>b</sup>abbreviations: lnSL = ln(sound length); logWF = log<sub>10</sub>(word frequency); PhonN = number of phonological neighbors; OrthN = number of orthographic neighbors; TTS = text-to-speech; $t_{MF}$ = multifractal nonlinearity; word# = word number in the presented sequence.
<sup>c</sup>boldfaced values indicate statistical significance at the alpha level of 0.05.

#### 3.1.1. Testing Hypothesis-1a: Multifractal nonlinearity sped the response in self-paced listening

Speech stimuli with greater multifractal nonlinearity evoked faster responses (*t*_MF_: *B* = –2.77×10^−5^, *p* < 0.01), although the difference was small, ranging from 0.90% to 2.74% as multifractal nonlinearity varied from 1^st^ to 3^rd^ quartiles (table 1). We found comparable effects of word frequency (logWF: *B* = –3.22×10^−2^, *p* < 0.0001) and phonological neighborhood size (PhonN: *B* = −3.97×10^−4^, *p* < 0.05) in speech perception. RT increased with logarithmically-scaled sound length (lnSL: *B* = 3.44×10^−1^, *p* < 0.0001): it takes longer to listen to longer sounds.

#### 3.1.2. Testing Hypothesis-1b: Multifractal nonlinearity deferred the slower response to text-to-speech synthesis—later than other psycholinguistic features did

Word-by-word RTs decreased at a power law (e.g., [72]; 1/word#: *B* = 1.02×10^0^, *p* < 0.001). Word-by-word RT increased linearly with each TTS word (TTS: *B* = −7.16×10^−2^, *p* < 0.0001; TTS×word#: *B* = 1.42×10^−4^, *p* < 0.0001; figure 5*a*). However, multifractal nonlinearity of speech sounds predicted a stronger power-law decay in RT in the TTS case (1/word#×TTS×*t*_MF_, *B* = 1.81×10^−3^, *p* < 0.01). Consequently, response to text-to-speech became significantly slower only much later for sounds with greater multifractal nonlinearity, with significant differences between RT for TTS and RT for human speech appearing on word #1140 for low *t*_MF_ (1^st^ quartile; thin gray vertical line figure 5*a*) and on word #1280 for high *t*_MF_ (3^rd^ quartile; thick gray vertical line in figure 5*a*). Greater multifractal nonlinearity led to comparable RTs for human speech and text-to-speech speech comparable for 140 words, or 12.28% longer than lower multifractality.

**Figure 5.**
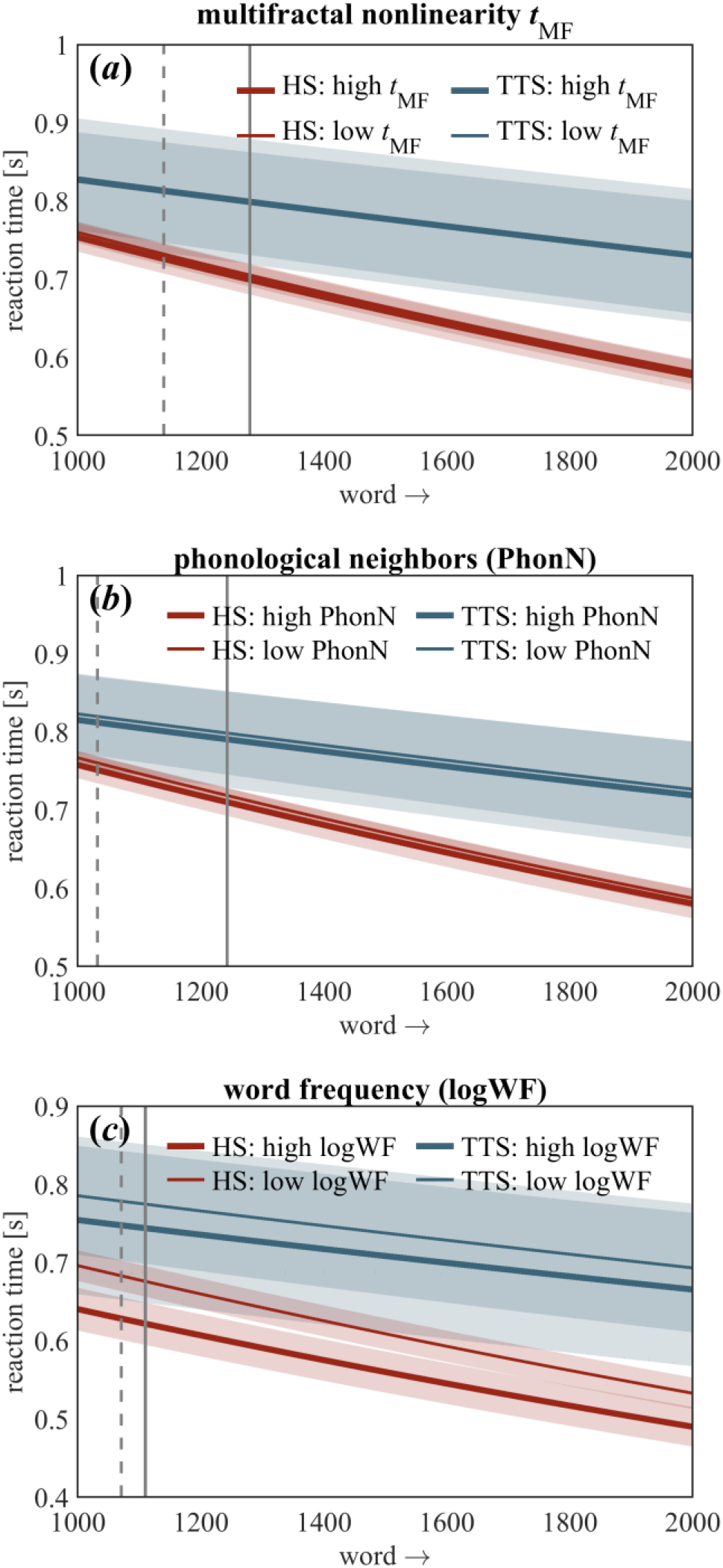
Model predictions for response time (section 3.1.2). Differences between human speech (HS) and text-to-speech (TTS) appear as marked by vertical dashed and solid lines for 1^st^ and 3^rd^ quartiles, respectively, of various predictors: words #1140 and #1280 for *t*_MF_ (*a*), words #1032 and #1243 for PhonN (*b*), and words #1071 and #1110 for logWF (*c*), respectively. Shaded bars indicate ±2*s.e.m.*

Effects of multifractal nonlinearity held above and beyond effects of standard psycholinguistic features. Phonological neighborhood yielded an even stronger power-law decay with text-to-speech (1/word#×TTS×PhonN: *B* = 8.21×10^−2^, *p* < 0.001); figure 5*b*), prolonging comparable RTs for text-to-speech and for human speech until about half way through the text. Frequency showed a comparable effect (TTS×logWF: *B* = 1.65×10^−2^, *p* < 0.0001; 1/word#×logWF: *B* = 2.84×10^−1^, *p* < 0.0001; figure 5*c*). Finally, orthographic neighborhood size exhibited no significant effect (*p* > 0.05), likely reflecting the auditory rather than the visual form of the task.

### 3.2. Testing Hypothesis 2: Model of cumulative correct responses on the 20-item two-part memory test

Story comprehension and word recognition for human speech (*mean* = 7.3 and 5.1 out of 10, *s.e.m.* = 0.45 and 0.46, respectively) were comparable to the same scores for text-to-speech (*mean* = 7.2 and 4.8 out of 10, *s.e.m.* = 0.39 and 0.66, respectively). Table 2 lists this model’s coefficients. Word recognition was easier (*B* = 2.99×10^−1^, *p* < 0.05), and text-to-speech synthesizers might have supported more effective learning with self-pacing (*B* = 4.25×10^0^, *p* < 0.01), whereas controlling the pacing of human speech might have been distracting.

**Table 2.**
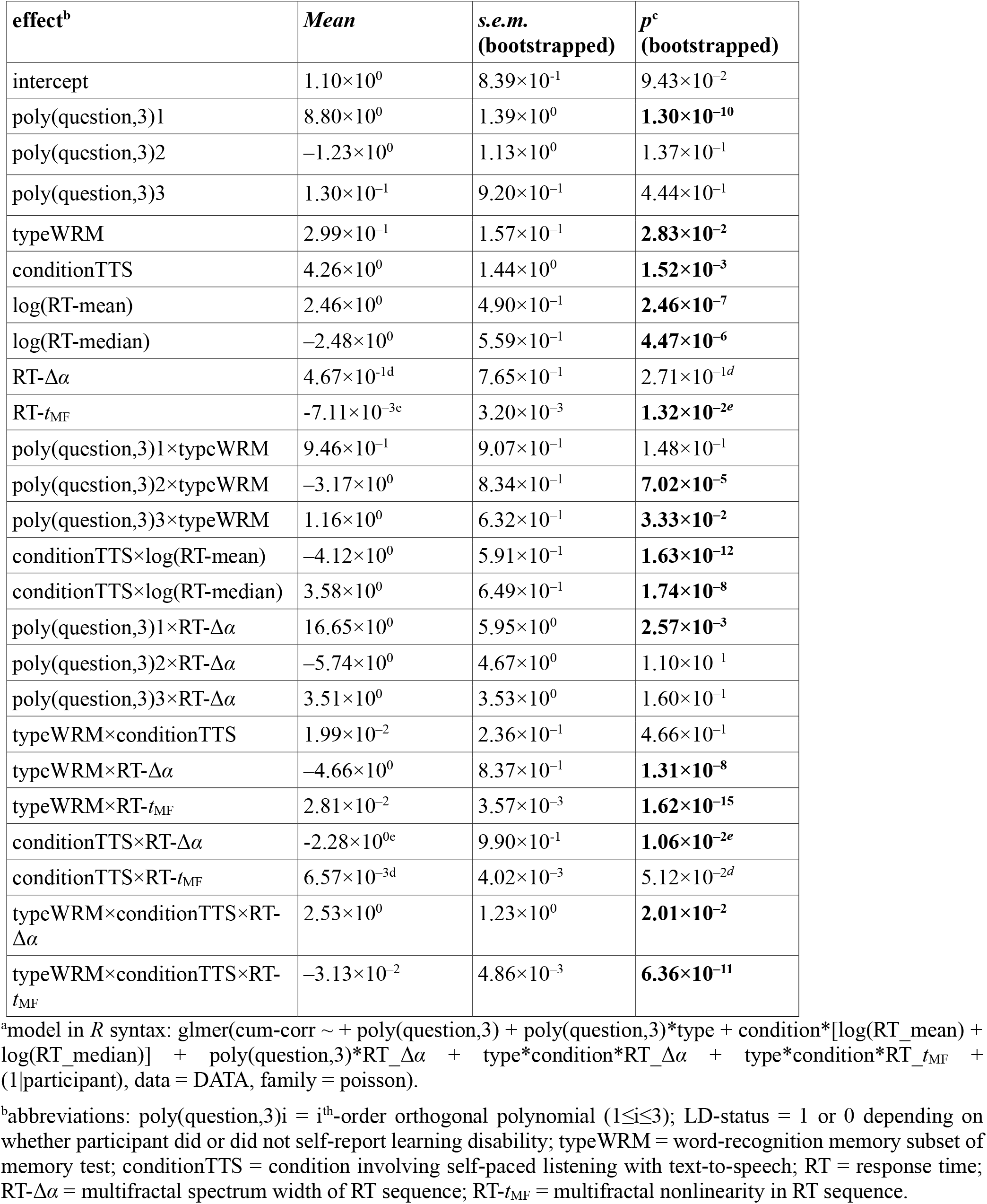
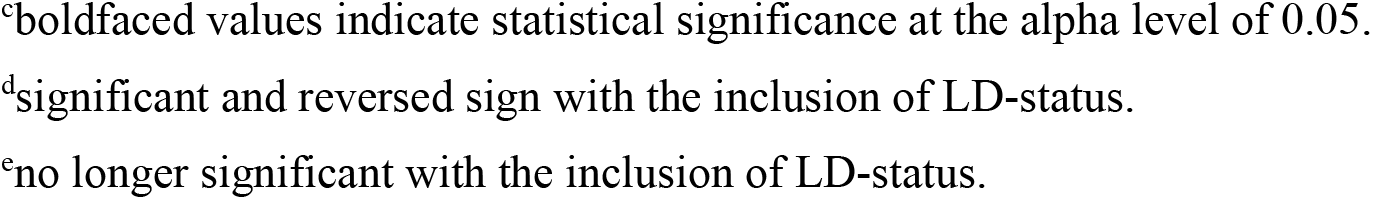
Coefficients of the Poisson regression of cumulative correct answers in the post-narrative test^a^ (section 3.2).

| effect<sup>b</sup> | *Mean* | *s.e.m.*<br>(bootstrapped) | *p*<sup>c</sup><br>(bootstrapped) |
| --- | --- | --- | --- |
| intercept | $1.10 \times 10^0$ | $8.39 \times 10^{-1}$ | $9.43 \times 10^{-2}$ |
| poly(question,3)1 | $8.80 \times 10^0$ | $1.39 \times 10^0$ | **$1.30 \times 10^{-10}$** |
| poly(question,3)2 | $-1.23 \times 10^0$ | $1.13 \times 10^0$ | $1.37 \times 10^{-1}$ |
| poly(question,3)3 | $1.30 \times 10^{-1}$ | $9.20 \times 10^{-1}$ | $4.44 \times 10^{-1}$ |
| typeWRM | $2.99 \times 10^{-1}$ | $1.57 \times 10^{-1}$ | **$2.83 \times 10^{-2}$** |
| conditionTTS | $4.26 \times 10^0$ | $1.44 \times 10^0$ | **$1.52 \times 10^{-3}$** |
| log(RT-mean) | $2.46 \times 10^0$ | $4.90 \times 10^{-1}$ | **$2.46 \times 10^{-7}$** |
| log(RT-median) | $-2.48 \times 10^0$ | $5.59 \times 10^{-1}$ | **$4.47 \times 10^{-6}$** |
| RT-$\Delta\alpha$ | $4.67 \times 10^{-1d}$ | $7.65 \times 10^{-1}$ | $2.71 \times 10^{-1d}$ |
| RT-$t_{MF}$ | $-7.11 \times 10^{-3e}$ | $3.20 \times 10^{-3}$ | **$1.32 \times 10^{-2e}$** |
| poly(question,3)1×typeWRM | $9.46 \times 10^{-1}$ | $9.07 \times 10^{-1}$ | $1.48 \times 10^{-1}$ |
| poly(question,3)2×typeWRM | $-3.17 \times 10^0$ | $8.34 \times 10^{-1}$ | **$7.02 \times 10^{-5}$** |
| poly(question,3)3×typeWRM | $1.16 \times 10^0$ | $6.32 \times 10^{-1}$ | **$3.33 \times 10^{-2}$** |
| conditionTTS×log(RT-mean) | $-4.12 \times 10^0$ | $5.91 \times 10^{-1}$ | **$1.63 \times 10^{-12}$** |
| conditionTTS×log(RT-median) | $3.58 \times 10^0$ | $6.49 \times 10^{-1}$ | **$1.74 \times 10^{-8}$** |
| poly(question,3)1×RT-$\Delta\alpha$ | $16.65 \times 10^0$ | $5.95 \times 10^0$ | **$2.57 \times 10^{-3}$** |
| poly(question,3)2×RT-$\Delta\alpha$ | $-5.74 \times 10^0$ | $4.67 \times 10^0$ | $1.10 \times 10^{-1}$ |
| poly(question,3)3×RT-$\Delta\alpha$ | $3.51 \times 10^0$ | $3.53 \times 10^0$ | $1.60 \times 10^{-1}$ |
| typeWRM×conditionTTS | $1.99 \times 10^{-2}$ | $2.36 \times 10^{-1}$ | $4.66 \times 10^{-1}$ |
| typeWRM×RT-$\Delta\alpha$ | $-4.66 \times 10^0$ | $8.37 \times 10^{-1}$ | **$1.31 \times 10^{-8}$** |
| typeWRM×RT-$t_{MF}$ | $2.81 \times 10^{-2}$ | $3.57 \times 10^{-3}$ | **$1.62 \times 10^{-15}$** |
| conditionTTS×RT-$\Delta\alpha$ | $-2.28 \times 10^{0e}$ | $9.90 \times 10^{-1}$ | **$1.06 \times 10^{-2e}$** |
| conditionTTS×RT-$t_{MF}$ | $6.57 \times 10^{-3d}$ | $4.02 \times 10^{-3}$ | $5.12 \times 10^{-2d}$ |
| typeWRM×conditionTTS×RT-$\Delta\alpha$ | $2.53 \times 10^0$ | $1.23 \times 10^0$ | **$2.01 \times 10^{-2}$** |
| typeWRM×conditionTTS×RT-$t_{MF}$ | $-3.13 \times 10^{-2}$ | $4.86 \times 10^{-3}$ | **$6.36 \times 10^{-11}$** |

<sup>a</sup>model in R syntax: glmer(cum-corr ~ + poly(question,3) + poly(question,3)\*type + condition\*[log(RT\_mean) + log(RT\_median)] + poly(question,3)\*RT\_Δα + type\*condition\*RT\_Δα + type\*condition\*RT\_t<sub>MF</sub> + (1|participant), data = DATA, family = poisson).
<sup>b</sup>abbreviations: poly(question,3)i = i<sup>th</sup>-order orthogonal polynomial (1≤i≤3); LD-status = 1 or 0 depending on whether participant did or did not self-report learning disability; typeWRM = word-recognition memory subset of memory test; conditionTTS = condition involving self-paced listening with text-to-speech; RT = response time; RT-Δα = multifractal spectrum width of RT sequence; RT-t<sub>MF</sub> = multifractal nonlinearity in RT sequence.
<sup>c</sup>boldfaced values indicate statistical significance at the alpha level of 0.05.
<sup>d</sup>significant and reversed sign with the inclusion of LD-status.
<sup>e</sup>no longer significant with the inclusion of LD-status.

Multifractality of RT from linear sources (i.e., Δ*α*) diminished memory performance only in word-recognition memory in the human-speech case (*B* = −4.66×10^0^, *p* < 0.0001; figure 6*b*). However, the text-to-speech manipulation showed smaller negative effects of multifractality from linear sources on both word-recognition memory and on narrative-comprehension memory (typeWRM×conditionTTS×RT-Δα: *B* = 2.53×10^0^, *p* < 0.05; conditionTTS×RT-Δα: *B* = −2.28×10^0^, *p* < 0.05; table 2). Multifractal nonlinearity reduced narrative-comprehension memory (B = −7.11×10^−3^, *p* < 0.05; figure 7*a*) but promoted word-recognition memory following self-paced listening to human speech (*B* = 2.81×10^−2^, *p* < 0.0001; figure 7*b*). From 1^st^ to 3^rd^ quartiles of RT-*t*_MF_, narrative-comprehension memory became 5% to 22% less accurate, and word-recognition memory became 25% to 150% more accurate (table 2). Memory performance on both tests decreased for participants with greater multifractal nonlinearity in text-to-speech condition only in word recognition (*B* = −3.13×10^−2^, *p* < 0.0001), but linear sources of multifractality promoted word-recognition memory.

**Figure 6.**
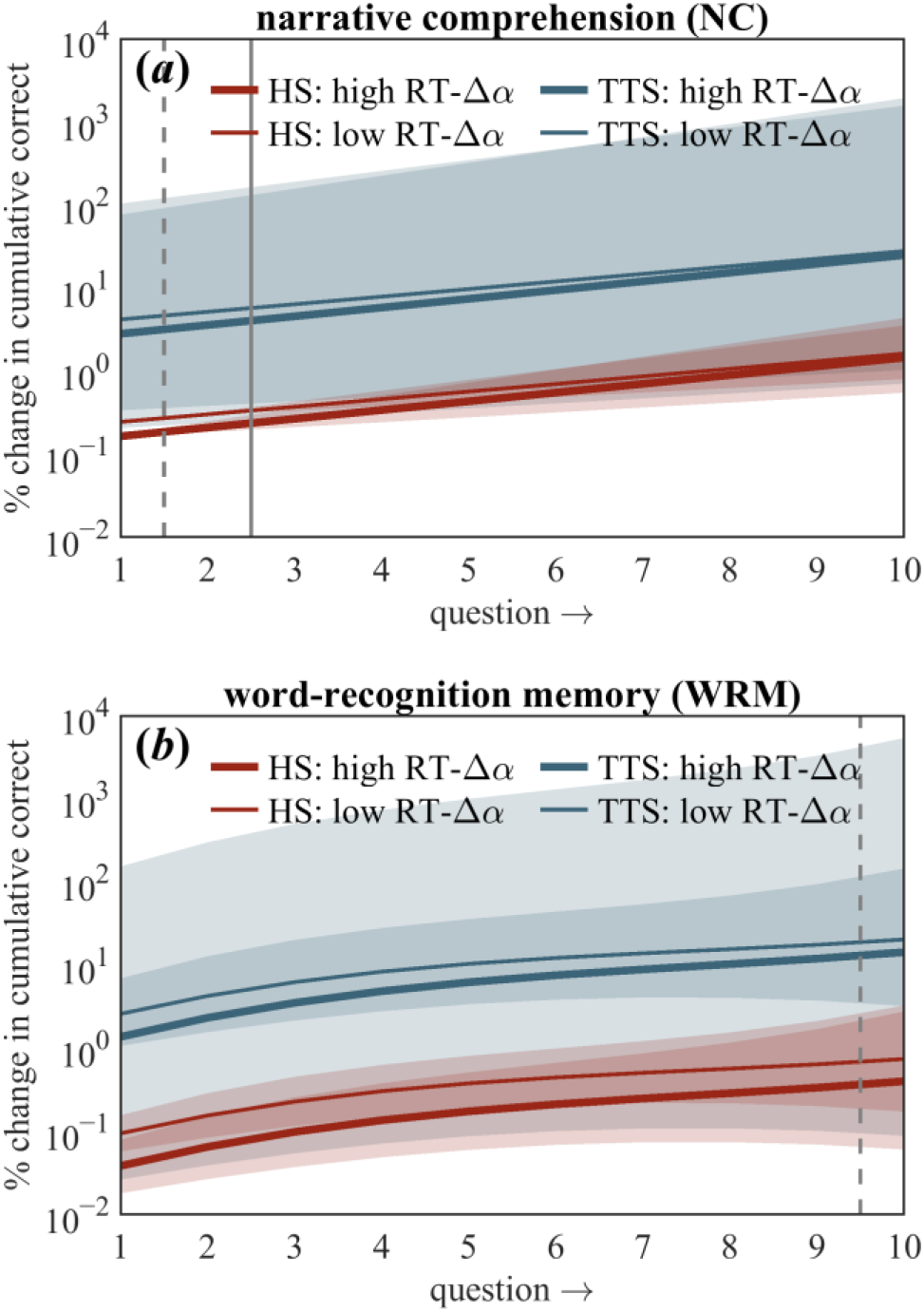
Model predictions for performance in narrative comprehension and word-recognition memory tests (section 3.2). Effect of RT-Δ*α* on % change in cumulative correct answers in narrative comprehension test (*a*) and in word-recognition memory test (*b*). Dashed and solid gray vertical lines indicate when low RT-Δ*α* and high RT-Δ*α*, respectively, stops predicting better test performance compared to low RT-Δ*α* and high RT-Δ*α*, respectively, in speech. Shaded bars indicate ±2*s.e.m.*

**Figure 7.**
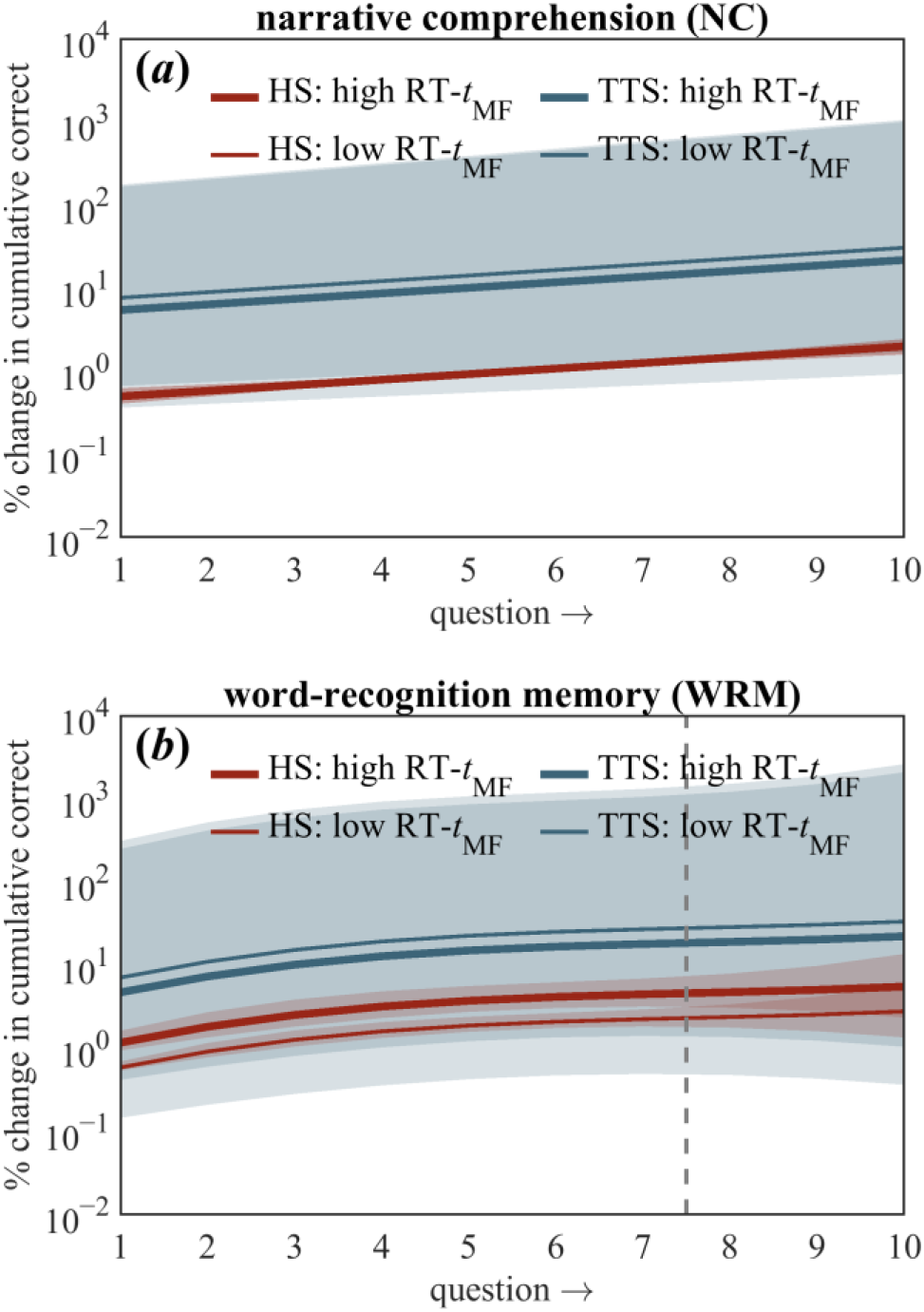
Model predictions for performance in narrative comprehension and word-recognition memory tests (section 3.2). Effect of RT-*t*_MF_ on % change in cumulative correct answers in narrative comprehension test (*a*) and in word-recognition memory test (*b*). Dashed vertical line indicates when low RT-*t*_MF_ stops predicting better test performance compared to low RT-*t*_MF_ in speech. Shaded bars indicate ±2*s.e.m.*

Over the course of each 10-question memory test, each correct response had the potential to prime more memory for the text. Correct responses increased at a linear relationship with question number (poly(Question, 3)1: *B* = 8.80×10^0^, *p* < 0.0001; figure 6). Multifractality from linear sources accentuated only the linear growth of correct responses with question number (i.e., Δ*α*; poly(Question,3)1×RT-Δ*α*: *B* = 16.65×10^0^, *p* < 0.01; figure 6). Multifractal nonlinearity did not show significant effects (table 2; figure 7). Correct responses in the word-recognition test increase across the 10 questions with a stronger cubic relation, including a negative quadratic relationship (poly(Question, 3)2×typeWRM: *B* = −3.17×10^0^, *p* < 0.0001; poly(Question, 3)3×typeWRM: *B* = 1.16×10^0^, *p* < 0.05; figures 6 and 7).

Post-text memory for the text showed a minor but significant decrease with longer RTs during the text for human speech (log(RT-mean): *B* = 2.46×10^0^, *p* < 0.0001; log(RT-median): *B* = −2.48×10^0^, *p* < 0.0001) and for text-to-speech (TTS×log(RT-mean): *B* = −4.12×10^0^, *p* < 0.0001; TTS×log(RT-median): *B* = 3.57×10^0^, *p* < 0.0001). Considering logarithm of mean and median RT together, longer (3^rd^ quartile) RTs predicted 4% more accurate memory performance than shorter RTs (1^st^ quartile) in human speech. The effect was smaller and reverse in the text-to-speech condition: 0.3% less accurate memory performance (table 2).

#### 3.2.1 Clarifying the Hypothesis 2 result: Controlling for LD-status effect indicates that multifractal nonlinearity only reduces narrative-comprehension in text-to-speech condition

We ran a second version of this model that included LD-status to control for the fact that readers with learning disabilities can benefit from self-pacing [31] and from their fluency using text-to-speech synthesis [78]. This predictor both improved model fit (*χ*^2^(1) = 4.65, *p* < 0.05) and demonstrated greater than 80% power in a 1000 bootstrapped resamplings of the data. This reanalysis confirmed that participants with learning disability did in fact remember more from the text (*B* = 3.75×10^−1^, *p* < 0.0001). Furthermore, including the LD-status predictor led to different estimates for main effects of RT multifractality and its interaction with condition. Specifically, including the LD-status predictor clarified that multifractal nonlinearity had no main effect, that the negative main effect actually belonged to multifractal linearity (i.e., RT-Δ*α*: *B* = −2.31×10^0^, *p* < 0.01), and that multifractal nonlinearity only predicted reduced memory in the text-to-speech condition (TTS×RT-*t*_MF_: *B* = −1.37×10^−2^, *p* < 0.05).

## 4. Discussion

We tested two hypotheses about the effects of multifractal nonlinearity *t*_MF_ in perception and memory for spoken language. Hypothesis-1 was that greater *t*_MF_ in speech stimuli would make speech perception more fluent, particularly in keeping responses text-to-speech synthesis as fluent as responses to human speech. Hypothesis-2 was that greater *t*_MF_ in the RT sequence in self-paced listening to a narrative would support better memory of the speech stimuli. Results supported Hypothesis-1 in full but Hypothesis-2 only in the word-recognition part of the memory test and not in the narrative-comprehension portion of the memory test. These findings carry implications not only for basic research into the movement basis for speech perception and memory for speech but also for the development of more effective text-to-speech synthesis for students with learning disabilities.

### 4.1. Multifractal nonlinearity promotes quicker response to speech stimuli and processing of text-to-speech synthesis later than traditional psycholinguistic features do

Multifractal nonlinearity of speech was associated with more fluent speech perception across a narrative. Whether the speaker was human or machine, the inevitable interactions across timescales as movement systems produce speech sounds (human speech and text-to-speech) promoted spoken-word perception in narrative sequence. Not only do articulatory motor processes matter to perception [8–11,80,81], their perceptual relevance goes deeper than linguistic distinctions (e.g., fricatives or stops and voice-onset times) and is rooted in more generic nonlinear dynamics with no currently recognized linguistic meaning [3,4]. The absence of any effects of multifractal spectrum width suggests that this aspect of speech perception depends on nonlinear and not linear relationships across timescales.

Multifractality exerted its role in supporting fluent processing of synthesized speech later than frequency and phonological neighborhoods, but longer than frequency supported. This result likely reflects that multifractal nonlinearity is a deeper attribute of acoustic waveform. Frequency and phonological neighborhood may support the fluent perception of less familiar speech early on because these are high-level features anchored in lexicality (i.e., word status). Meanwhile, multifractal nonlinearity is not specific to words and inheres in the heterogeneous movements producing speech sounds. Perception resorts most clearly to this multifractal movement foundation when linguistic support for word recognition falters.

### 4.2. Multifractal nonlinearity in self-paced listening to human speech promoted word-recognition memory, and multifractality from linear sources predicted diminished memory on average but increased memory performance as testing continued

Participant’s ability to remember the speech stimuli exhibited a more complex relationship between movement systems across the speaker-listener dyad. The two movement systems generating speech (i.e., natural human speech vs. computer-simulated analogue) cultivated distinct aspects of the multifractal structure of self-pacing responses: comprehension of more familiar human speech benefited from nonlinear aspects of multifractal self-pacing behaviors (i.e., *t*_MF_), and comprehension of less familiar text-to-speech benefited from linear aspects (i.e., Δ*α*). Self-paced listening to speech presented word by word may have emphasized individual words at the expense of attention to the narrative’s discourse-level aspects. Also, it is possible that including seven true and three false items in the comprehension test reduced variability in responses (e.g., by cuing more memory of the text).

Multifractal nonlinearity appeared at first to diminish narrative-comprehension memory, but controlling for learning-disability advantages with self-paced listening showed instead that multifractal nonlinearity only diminished narrative-comprehension memory in the text-to-speech condition. This point is intriguing because multifractality nonlinearity refers to the interaction of short-range (e.g., word-by-word) scales of activity with long-range (e.g., discourse/narrative) scales of activity. So, it could be that self-paced listening behaviors with this cross-scale interactivity makes maladaptive alignment with text-to-speech synthesis that only produces speech at word-by-word scales.

## 5. Conclusions

The present work is the latest step in the new enterprise of modeling multifractal effects supporting spoken language use. It addresses current calls in the literature for a straightforward multifractal extension of the notion that a speaker-listener dyad depends on ‘complexity matching’ between the respective movement systems [56–59]. First, the multifractal evidence of nonlinear interactions across timescales predicts the fluency of speech perception. Second, self-pacing movements that enabled listeners to control the presentation of speech sounds constituted a rich exploratory process. Multifractal nonlinearity of this exploration supports some but not yet all aspects of memory for perceived spoken language. Nonetheless, the present findings suggest an effect of multifractal structure in the both speech production and listener’s exploratory movements in speech perception.

Self-paced listening is not ecologically valid for speech perception “in the wild” (e.g., a listener listening to a speaker producing words at the speaker’s own preferred pace). However, future work could examine whether the multifractal structure of postural or head sway could moderate speech perception as it has already been shown to moderate visual and haptic perception (e.g., [47,48,51,52,82]). Even admitting the contrived status of self-paced listening, the present results validate our idea that speech perception and memory reflects an alignment (or not) between the structures of the speaker’s and the listener’s movements. Speech memory reflected a compromise between the speaker’s movement system and the listener’s actions’ multifractality. The self-pacing paradigm prompting the listener to act to get the next word coaxes some movement whose geometry is also multifractal. Different movement systems can cultivate specifically different (i.e., linear or nonlinear) aspects of the multifractality in self-paced timing. A more familiar speaker (i.e., human speech) sooner leads to multifractal nonlinearity to support speech memory. In contrast, the less familiar speaker (i.e., text-to-speech [14]) sooner leads to linear sources of multifractality to support speech memory.

Future work could investigate aspects of human or text-to-speech voices influencing relative linear or nonlinear effects of multifractality on perception and subsequent memory for speech. For instance, if multifractality influences familiarity of speech, then it could hence enhance text-to-speech synthesis as a learning tool. The present evidence that multifractality of speech movements is essential for speech perception is silent on whether multifractality of speech movements have consequences for memory. Unambiguous comparison of multifractality across dyads was precluded by the radically different data formats between speaker and listener recordings (multiple 44.1 kHz digital recordings vs. single sequences of 2,026 inter-key press intervals). Whether this alignment rests firmly on multifractality could be tested in a ‘shadowing’ paradigm (i.e., listeners repeating immediately each spoken word, e.g., [83]). It could be beneficial to test how greater familiarity with a particular voice influences the appearance and role of multifractality. Another important point for future work is that words appeared in the narrative context, so it may be worth extending this work to a fuller accounting of RTs not just for words but also for phrases, sentences, paragraphs, etc. [71].

## Supporting information

Self-paced individual-word response times

Individual memory-test response data

Memory test

## Supplementary materials

**Supplementary Note S1.** Reading comprehension test given to each participant.

**Supplementary Dataset S1.** Data modeled in the Poisson regression of self-paced word RT (section 3.1).

**Supplementary Dataset S2.** Data modeled in the Poisson regression of cumulative correct answers in the post-narrative test (section 3.2).

## Notes

### Competing Interest Statement

The authors have declared no competing interest.

### Summary of Updates

Fixing typos

