## Supplementary material for "Perceiving and remembering speech depend on multifractal nonlinearity in movements producing and exploring speech": Memory test

### Reading Comprehension Test

Please answer the following questions based on whether this information is true or false based on the passage you just heard. Please circle either true or false.

1. **(T/F)** If the books were scanned, they would have been available for free at terminals in local libraries.
2. **(T/F)** On the scanned copies, you would not be able to annotate the books.
3. **(T/F)** After the books were scanned, the books would become like webpages.
4. **(T/F)** It had never been previously imagined amassing a universal library before Google Books made it a reality.
5. **(T/F)** Many scholars, archivists and libraries were not in support of the digital book sharing.
6. **(T/F)** Google was trying to secretly scan every book in the world.
7. **(T/F)** To better the search results book results would be sorted by date.
8. **(T/F)** The stations that digitally converted the books initially took photographs of each page.
9. **(T/F)** Google fell a hundred-million books short of scanning all 129,864,880 books in the world.
10. **(T/F)** Google had no legal issues with copyright infringement in copying the books because they always asked for permission before scanning.

Please circle either true or false as to whether or not you heard the following words in the passage.

1. **(T/F)** Precisely
2. **(T/F)** Millennia
3. **(T/F)** Watershed
4. **(T/F)** Confusion
5. **(T/F)** Peppered
6. **(T/F)** Metronome
7. **(T/F)** Revolutionary
8. **(T/F)** Algorithms
9. **(T/F)** Boondoggle
10. **(T/F)** Profit

Please write a brief summary of the passage in three sentences or less.[a]

____________________________________________

____________________________________________

____________________________________________

[a]This summary question was included to confirm that participants paid attention to the gist, and we did not quantify their open-ended responses.
